# Development of a Genetically Encoded Live-Cell Iron Mapping System

**DOI:** 10.1101/2025.02.25.639876

**Authors:** Ali Akyol, Şeyma Çimen, Benjamin Gottschalk, Amy Barton Alston, Reinaldo Digigow, Beat Flühmann, Emrah Eroğlu, Wolfgang F. Graier, Roland Malli

**Affiliations:** Department of Molecular Biology and Biochemistry, Gottfried Schatz Research Center, Medical University of Graz, Austria; Regenerative and Restorative Medicine Research Center (REMER), Research Institute for Health Sciences and Technologies (SABITA), Istanbul Medipol University, Istanbul, Turkey; CSL Vifor Ltd., Redwood City, CA, USA; CSL Vifor Ltd., Glattbrugg, Switzerland; CF Bioimaging, Center of Medical Research, Medical University of Graz, Austria; BioTechMed Graz, Austria

## Abstract

Any disturbances in cellular iron regulation can lead to dysfunctions and various diseases. However, our current understanding of iron dynamics in living cells is hampered by the lack of high-resolution iron reporter systems capable of monitoring the iron status of individual cells dynamically. To address this challenge, we have developed the Live-Cell Iron Mapping System (LIMS), which consists of two distinct genetically encoded fluorescent iron reporter systems: the rFeRepS and IronFist. The rFeRepS (ratiometric Iron Reporter Systems) code for two genetically encoded constructs that translate the iron status of individual cells into a ratiometric fluorescent signal. The ratio signal reflects the cellular iron status, based on endogenous iron-responsive proteins. A high rFeRepS ratio indicates ferritin-based cellular iron storage, while a low ratio implies transferrin receptor (TfR)-based active cellular iron uptake. The IronFist (Iron Fluorescent Indicators based on Stability and Translation) is derived from the hemerythrin-like domain (Hr) of the F-box and leucine-rich protein 5 (FXBL5), which undergoes a structural rearrangement upon binding to ferrous iron, preventing its ubiquitination and subsequent degradation. In the IronFist design, we fused either the blue or green fluorescent protein (FP) variant, mTagBFP2 or mNeonGreen, to the C-terminus of Hr and co-expressed mCherry via a ribosomal skipping sequence as a reference protein for normalization. Thus, IronFist dynamically translates fluctuations of the labile iron pool into a ratiometric fluorescent signal. We demonstrated the functionality of both iron reporter systems by treating cells with different iron formulations including iron carbohydrate nanoparticles. Our results show clear cell-to-cell variations in the response to high iron treatments.

## Physiological Mechanisms Underpinning FeReps and IronFist: Core Components of the Live-Cell Iron Mapping System (LIMS)

The **LIMS** combines at least three distinct signals to monitor cellular iron status dynamically. **Signal 1** indicates elevated iron levels, **Signal 2** reflects reduced iron presence, and **Signal 3** specifically tracks changes in the labile iron pool (LIP), providing rapid and reversible feedback on acute iron fluctuations. While Signals 1 and 2 provide a long-term ratio metric overview of cellular iron status, Signal 3 enables real-time monitoring of labile iron dynamics (**Figure 1**).

**Figure 1.**
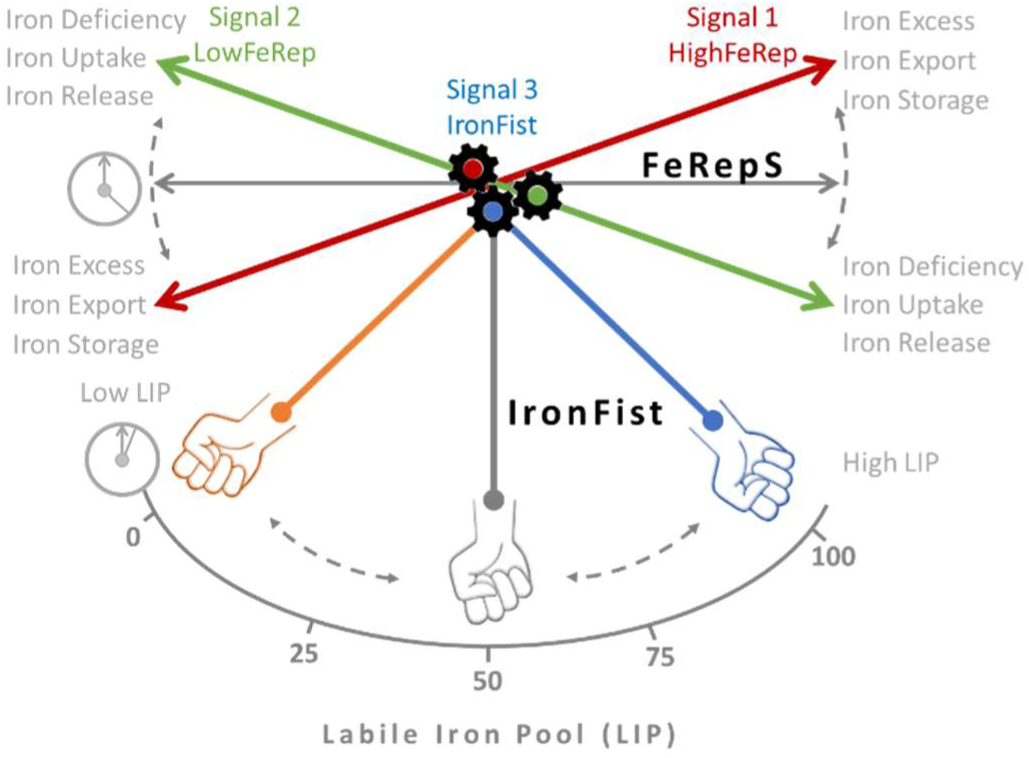
Assessing cellular iron status using a genetically encoded iron-mapping system. Schematic representation illustrating the interaction between FeRepS and IronFist in monitoring cellular iron levels. FeRepS comprises two signals (Signal 1 and Signal 2) that shift in response to changes in iron availability, indicating either iron excess (red) or iron deficiency (green). Simultaneously, IronFist dynamically responds to alterations in the labile iron pool (LIP) with rapid kinetics, providing real-time insights into changes in cellular iron(II) levels.

The three components and their respective signals (Signal1, Signal2, and Signal3) can be used independently; however, their combination provides enhanced insights into the dynamic cellular iron status — encompassing iron uptake, release, export, and storage — and facilitates a more comprehensive interpretation of whether an individual cell is in a state of iron deficiency, iron overload, or balanced iron supply.

The FeReps and IronFist systems are integral to the Live-Cell Iron Mapping System (LIMS), visualizing the precise regulation of cellular iron homeostasis. The regulation of the cellular iron status is carefully controlled by the expression of key proteins—most notably, the transferrin receptor (TfR) and ferritin (Fr)—to maintain optimal iron levels within cells (Pantopoulos, 2004; Zhou & Tan, 2017).

As shown in **Figure 2**, this regulatory mechanism relies on iron-responsive elements (IREs) and iron regulatory proteins (IRPs), including aconitase 1 (Aco1, also known as IRP1) and aconitase 2 (Aco2, also called IRP2)(Henderson et al., 2005). A unique feature of IRP2 is its lack of an iron-sulfur cluster (ISC) binding domain, which allows it to modulate ferritin synthesis and stabilize TfR mRNA differently from IRP1. The stability of IRP2 is regulated by FBXL5, a protein that targets IRP2 for proteasomal degradation in the presence of Fe²⁺ (Figure 2). In low-iron conditions, FBXL5 is itself degraded through polyubiquitination, leading to the accumulation of IRP2 and promoting TfR expression for increased iron uptake (Gárate & Núñez, 2000).

**Figure 2.**
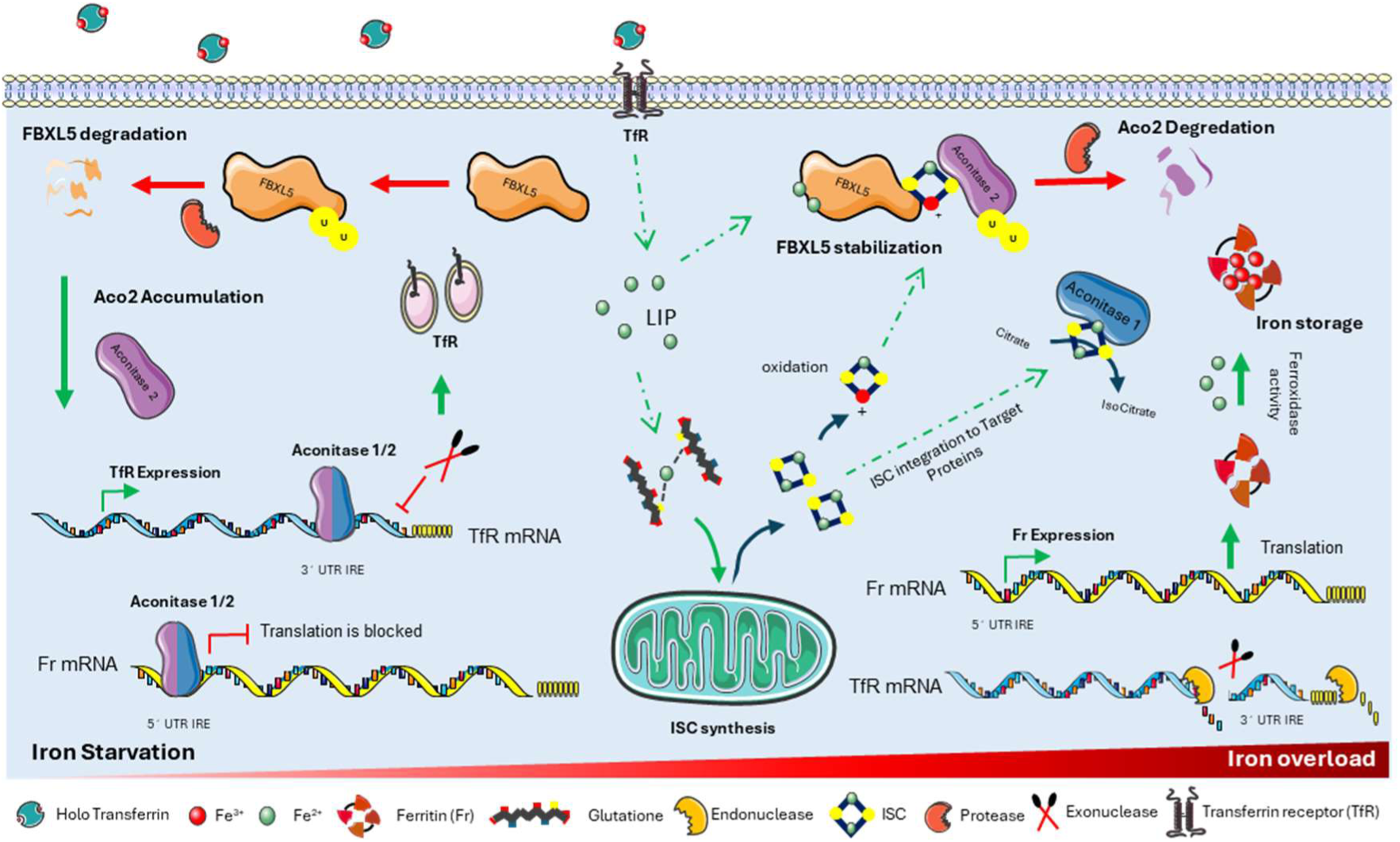
Key mechanisms controlling the cellular iron status. Schematic representation of the role of FBXL5 in maintaining cellular iron homeostasis and its interaction with the IRE-IRP system. Iron uptake via transferrin through the TfR pathway leads to an increase in the labile iron pool (LIP). The lysosomal reduction and release of Fe²⁺ are not shown here. An increase in LIP stabilizes FBXL5, preventing its degradation. The stabilized FBXL5 then binds to Aco2, leading to its degradation (right side of the figure). As a result, ferritin (Fr) expression increases to store excess iron, while transferrin receptor (TfR) expression is reduced, thereby limiting further iron uptake.

As cellular iron (II) levels (labile iron pool, or LIP) rise through extracellular intake or ferritin recycling (ferritinophagy),(Ohshima et al., 2022) IRP2 is progressively downregulated, with FBXL5 stabilization marking sufficient iron availability. We think that oxidized ISC signals that ISC needs are met, enabling both IRPs to detach from their mRNAs, and promoting ferritin synthesis to lower excess LIP(Torti & Torti, 2002) **Figure 2** illustrates this feedback loop, central to iron regulation in the LIMS framework.

## RESULTS and Discussions

### The rFeRepS (ratiometric Iron Reporter Systems): Working principle and construct design

The rFeRepS are genetically encoded constructs that translate the iron status of individual cells into a ratio metric fluorescent signal, where the ratio of e.g. *F*_mCherry_/*F*_mNeonGreen_ reflects cellular iron homeostasis, based on endogenous iron-responsive proteins (**Figure 3A**); a high ratio suggests ferritin-based cellular iron storage, while a low ratio implies transferrin receptor (TfR)-based active cellular iron uptake. To translate the iron status of living cells into a (optical) ratio metric readout, we introduce the genetically encoded rFeRepS (ratio metric Iron Reporter Systems). These systems convert alterations in cellular iron status into a detectable ratio metric signal by the opposing expression changes in at least two distinct proteins that represent or generate two distinct signals — such as distinct fluorescent protein variants of different colors (**Figure 3**). The rFeRepS displays the elevation of one signal concomitant with heightened iron supply while simultaneously suppressing another signal (**Figure 3**). Conversely, in conditions of cellular iron deficiency, the inverse occurs: the suppressed signal ascends while the initially elevated signal diminishes.

**Figure 3.**
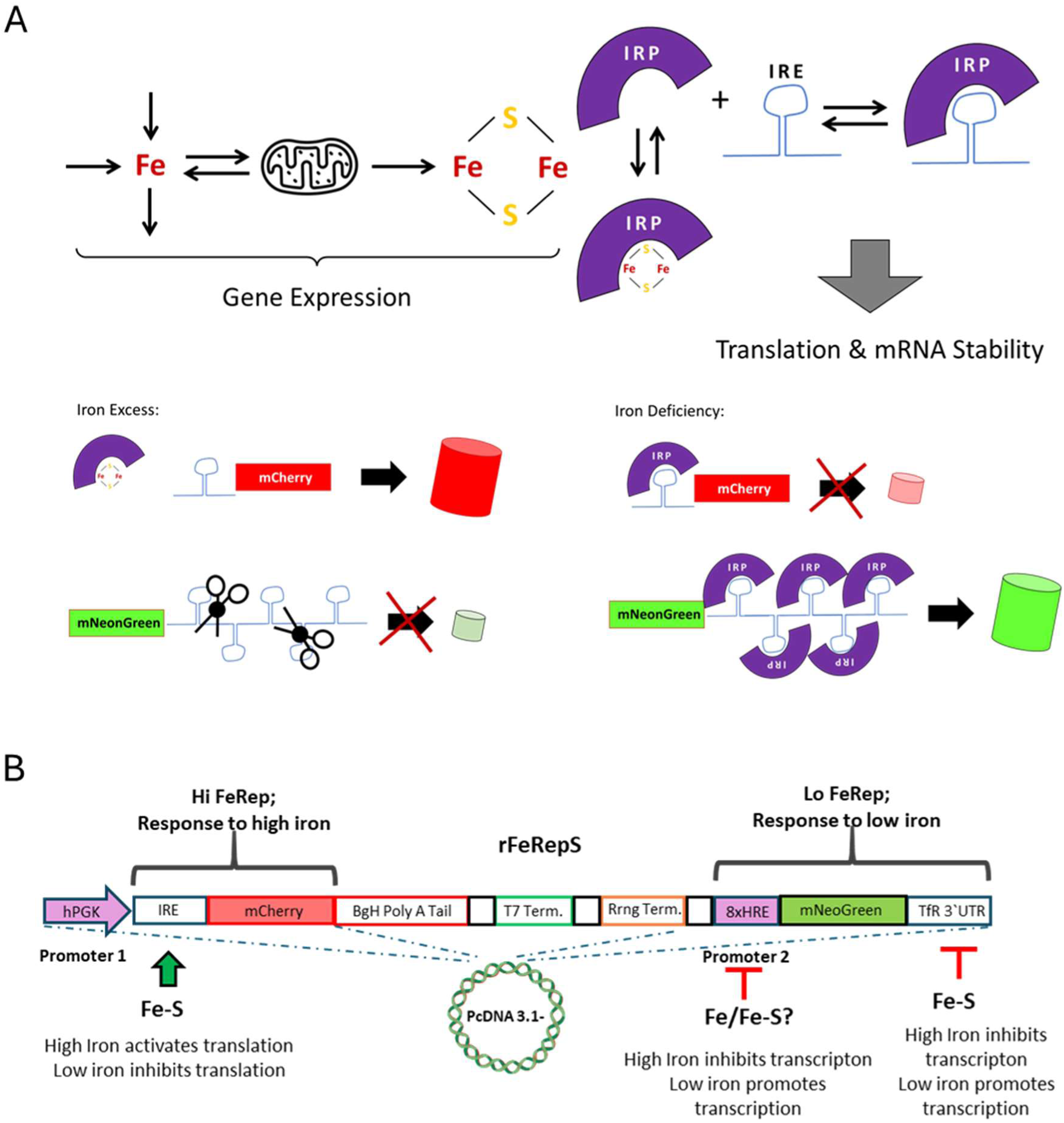
Principle and Design of Ratio Metric Iron Reporter Systems (rFeRepS). **A**) *Upper panel:* Iron within a cell is utilized for iron-sulfur cluster (ISC) biosynthesis, a process for which mitochondria are key organelles. ISC export from mitochondria allows integration into cytosolic proteins, such as iron regulatory proteins (IRPs), which control protein expression by interacting with iron response elements (IREs) on respective mRNAs. *Lower panel:* We have harnessed this mechanism to create a system that converts changes in cellular iron levels into a detectable red-green ratio metric fluorescent signal using a genetically encoded approach. **B**) *rFeRepS DNA Structure:* We designed a plasmid derived from the PcDNA3.1-vector, incorporating iron response elements (IREs) on mRNAs regulated by endogenous iron-responsive proteins (IRPs). The system enables the opposing expression of two distinct fluorescent proteins, mCherry and mNeonGreen, depending on cellular iron levels. The hPGK promoter drives expression of the Hi FeRep mRNA, while the BgH Poly A Tail, T7 terminator, and Rrng terminator, all derived from the original PcDNA3.1-vector, terminate Hi FeRep expression. Lo FeRep: The 8xHRE (hypoxia-responsive element) promoter region from the TfR gene controls mNeonGreen expression under low iron conditions. Additionally, the stability of the mNeonGreen mRNA is regulated by five IRE repeats in the 3’ UTR, reflecting the sequence of the TfR 3’ UTR. The original BgH Poly A Tail and terminator sequences are included after the TfR 3’ UTR (not shown).

To enable the expression of mCherry and mNeonGreen based on cellular iron status, we designed and constructed a modified PcDNA3.1-vector (**Figure 3B**). This vector allows for the independent expression of each fluorescent protein variant in response to iron levels within the cell. Specifically, mCherry expression mimics the expression pattern of Fr, increasing when cellular iron levels are high, while mNeonGreen expression reflects the behaviour of the TfR, increasing when iron levels are low. We replaced the CMV promoter in the standard PcDNA3.1-vector with the moderate-strength human phosphoglycerate kinase (hPGK) promoter to drive mCherry mRNA expression. This lower expression level helps maintain endogenous regulation by Aco1 and Aco2, ensuring that the mCherry signal mainly reflects Fr expression. When Aco1 and Aco2 detect high iron levels, they dissociate from the mRNA, enabling translation of the red fluorescent protein variant. This results in the "Signal 1" (Figure 1), referred to as "**Hi FeRep**" (Figure 3B). To terminate mCherry expression and enable independent expression of a second mRNA, we inserted an additional polyA tail and terminator signals from the original PcDNA3.1 vector at the end of Hi FeRep. The **Lo FeRep** system is controlled by the 8xHRE (hypoxia-responsive elements) promoter region from the TfR1 gene (Johansson et al., 2017), which is sensitive to low iron levels. This promoter drives mNeonGreen expression when cellular iron is low, reflecting the need for increased TfR expression to enhance iron uptake from the extracellular space. Additionally, the stability of the mNeonGreen mRNA is regulated by five IRE repeats in the 3’ UTR, which control translation under low iron conditions. Together, the iron-sensitive 8xHRE promoter and the IRE repeats in the 3’ UTR ensure that mNeonGreen expression is activated when iron levels fall, signaling the cell’s need for more iron.

### Hi FeRep dynamically reports changes in the cellular iron content

To investigate the dynamic response of the Hi FeRep construct to changes in cellular iron levels, we performed a series of experiments in HeLa cells, comparing the Hi FeRep (**Figure 4, left panel**) with the iron-insensitive control construct (Hi FeRep-IRE, **Figure 4, right panel**). Fluorescence intensity in cells expressing each construct was monitored over time under three different treatment conditions to evaluate iron-responsive expression kinetics (**Figure 4**).

**Figure 4.**
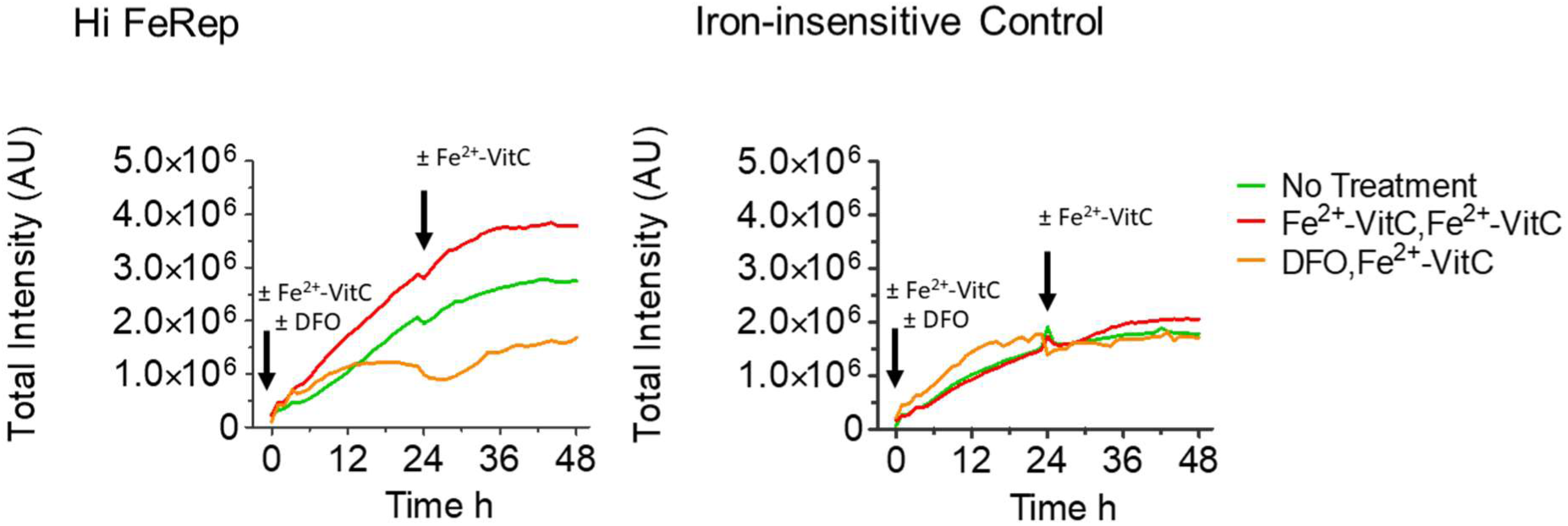
Dynamic response of Hela cells expressing Hi FeRep or iron insensitive control construct (Hi FeRep -IRE) **A)** Average curves (n=3 wells) showing total fluorescence intensity of HeLa cells expressing Hi FeRep, monitored over time. **B)** Total fluorescence intensity of HeLa cells expressing the iron-insensitive control construct i.e. Hi FeRep-IRE, monitored over time (average curves, n=3 wells). In both panels, the red curves represent cells treated twice (0h and 24 h) with Fe²⁺ combined with Vitamin C (150 µM FeSO₄ and 250 µM Vitamin C), to increase intracellular iron levels. The orange curves show cells pre-treated with 100 µM DFO (deferoxamine) at time zero; 24 hours later, the media was replaced with Fe²⁺ combined with Vitamin C to restore iron levels. The green curves represent untreated control cells, which received no additional iron or chelator treatment.

In the first condition, cells were treated with Fe²⁺ combined with Vitamin C to elevate intracellular iron levels and assess the construct’s response to iron enrichment. Cells exposed to Fe^2+^/Vitamin C developed Hi FeRep fluorescence intensity at a faster rate than untreated controls and DFO-treated cells (**Figure 4, left panel**), indicating that Hi FeRep expression is responsive to elevated iron levels. In contrast, cells expressing the iron-insensitive construct (Hi FeRep-IRE) showed no significant changes in fluorescence intensity or expression kinetics across treatments, demonstrating its lack of responsiveness to iron modulation (**Figure 4, right panel**). In the second condition, cells were initially treated with DFO to deplete cellular iron levels; 24 hours later, the media was replaced with Fe²⁺ and Vitamin C, allowing us to observe the construct’s response to iron reintroduction following an iron-depleted state. Cells treated with DFO showed a much slower increase in mCherry fluorescence over time compared to control cells and cells treated with the Fe(II) salt. The re addition of iron slightly restored the fluorescent intensity (**Figure 4, left panel**). In the third condition, untreated control cells were imaged over time to get a baseline fluorescence intensity for comparison (**Figure 4**).

### ELT treatment significantly increases the number of cells expressing Lo FeRep

To assess the performance of Lo FeRep, we expressed the construct in HeLa cells and treated them with eltrombopag (ELT) (**Figure 5**). Under control conditions, only 6.8% of cells showed a detectable Lo FeRep signal, indicating that a small proportion of HeLa cells were experiencing iron deficiency (**Figure 5B**). Treatment with the iron chelator ELT induced cell death and did not increase mNeonGreen fluorescence intensity in individual cells (data not shown). However, in response to ELT, we observed a significant increase in the number of cells positive for mNeonGreen (**Figure 5A and B**). These results suggest that the increase in Lo FeRep signal observed at the 24-hour endpoint likely represents an underestimation of iron deficiency. The cytotoxic effect of ELT may lead to the loss of Lo FeRep-positive cells, particularly those suffering from severe iron deficiency due to ELT treatment.

**Figure 5.**
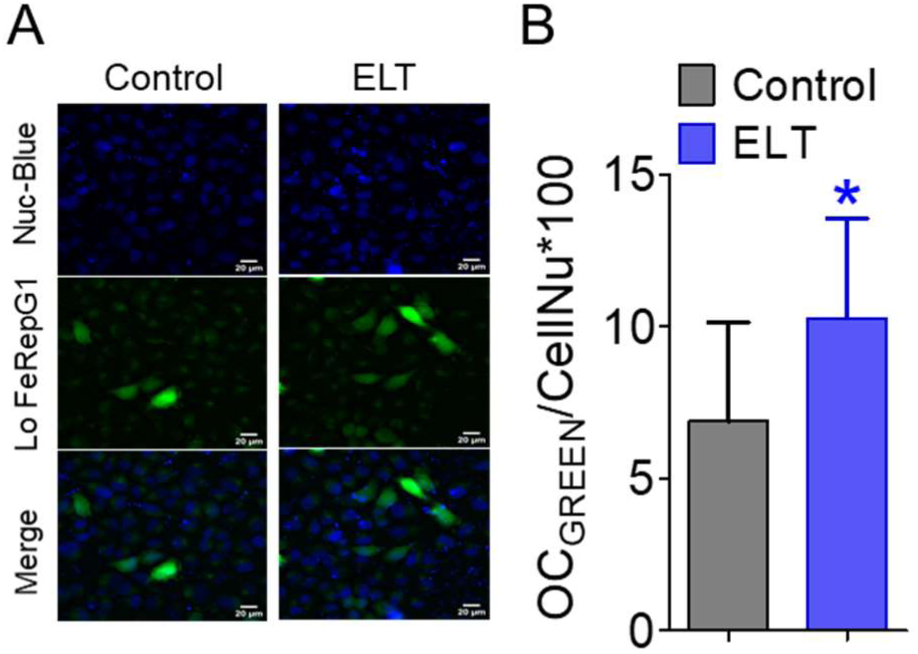
Response of Lo FeRep to ELT treatment in HeLa cells. **A**) Representative images of HeLa cells expressing Lo FeRep, with nuclei stained in blue to count the number of cells per imaging area. The green channel shows the Lo FeRep signal, and the merged image combines both channels. **B**) Bar graph showing the average percentage of Lo FeRep-positive cells, with error bars representing DATA=mean ±SD. The grey bars (n=3 wells) represent control conditions, while the blue bars (n=3 wells) show cells treated with ELT. * Indicate statistically significant differences (p < 0.05) between control and ELT-treated conditions.

### Assessment of cellular iron status using rFeRepS in HeLa cells

To explore the iron-sensing capability of rFeRepS, a construct designed a plasmid to express mCherry and mNeonGreen in an iron-dependent manner (**Figure 3B**), we transiently transfected HeLa cells and analyzed fluorescence signals post-transfection. We generated ratiometric images based on mCherry/mNeonGreen fluorescence intensity (**Figure 6A, right image**). Across multiple imaging areas, we observed a heterogeneous population of cells: some displayed a low mCherry/mNeonGreen ratio, indicative of low mCherry but high mNeonGreen fluorescence, while others showed a high mCherry/mNeonGreen ratio, with high mCherry and low mNeonGreen fluorescence. According to the rFeRepS functionality, a low red/green fluorescence ratio corresponds to low intracellular iron supply, while a high ratio suggests higher, potentially excessive iron accumulation.

**Figure 6.**
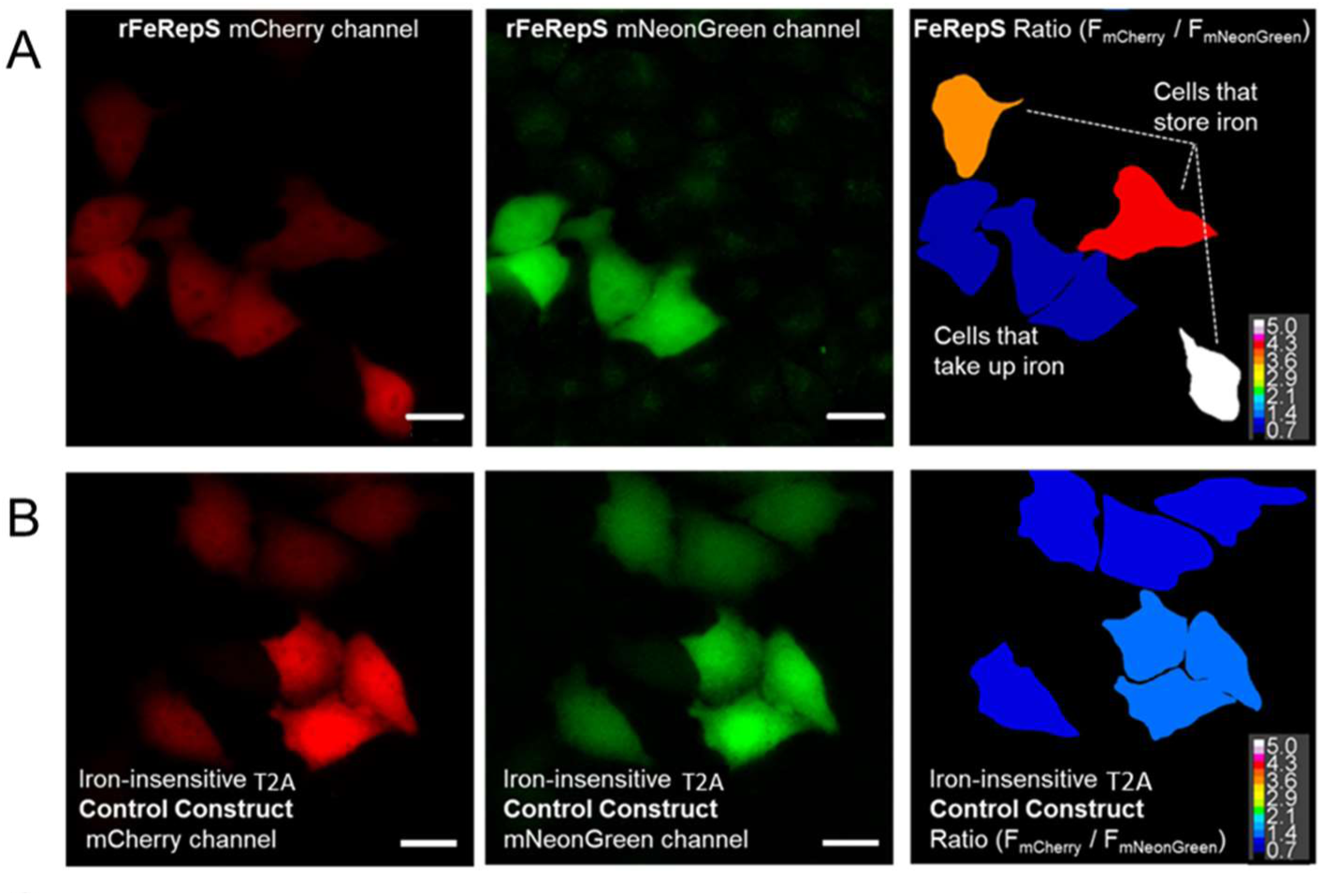
Evaluation of rFeRepS iron-sensing capabilities in HeLa cells. **A)** Representative images of HeLa cells transiently expressing rFeRepS. The left image shows mCherry fluorescence, the middle image shows mNeonGreen fluorescence, and the right image displays the mCherry/mNeonGreen ratio image, highlighting cells with varying iron levels. **B)** Representative images of HeLa cells expressing the control construct, designed to produce mCherry and mNeonGreen independently of iron levels. In these control cells, fluorescence intensity ratios remain consistent across the cell population, confirming that variations in Panel A are due to iron-sensing elements within rFeRepS. C) Schematic diagram of the control construct design. Both mCherry and mNeonGreen are expressed under the hPKG promoter with a T2A ribosomal skipping sequence, ensuring equal and iron-independent expression of both fluorescent proteins.

As a control, we expressed both fluorescent proteins in HeLa cells, without iron-responsive elements but under the same promoter strength. In this setup, all cells exhibited a consistent red/green ratio (**Figure 6B and C**), confirming that variability in fluorescence ratios observed with rFeRepS is due to differences in cellular iron levels rather than expression variability.

Based on these findings, we hypothesize that rFeRepS reliably indicates cellular iron status. To further validate this, we treated the transfected cells with Fe(II) sulfate/Vitamin C, iron sucrose nanoparticles (IS), and ELT (Vlachodimitropoulou et al., 2017), to assess whether these manipulations would shift the fluorescence ratios in accordance with altered cellular iron levels.

To examine the distribution of the mCherry/mNeonGreen ratio within the HeLa cell population expressing rFeRepS and investigate any potential correlation with expression levels, we plotted the red/green ratio values (x-axis) against the total fluorescence intensity (summed mCherry and mNeonGreen fluorescence per cell) in an XY scatter plot under control (untreated) conditions (**Figure 7A, left upper panel, B and C**). This analysis revealed substantial heterogeneity in red/green ratio values across the cell population, with no clear correlation between the ratio values and overall expression levels. We observed cells with both high and low ratio values, indicative of varying iron states, regardless of expression level. This suggests that cells with either low or high iron supply are present across a range of expression intensities.

**Figure 7.**
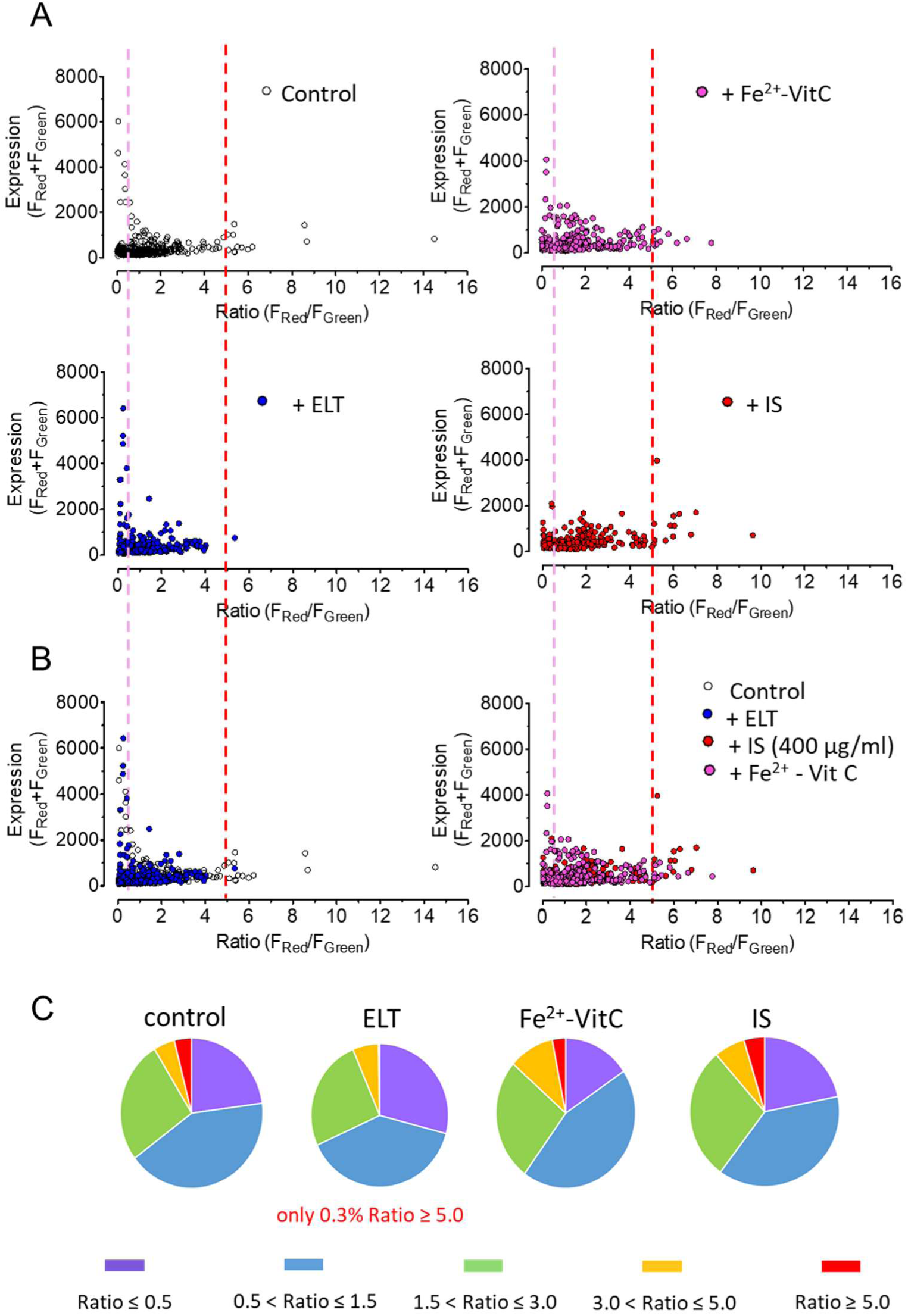
Distribution of mCherry/mNeonGreen Ratios in HeLa Cells Under Different Treatment Conditions. **A)** XY plots showing the distribution of mCherry/mNeonGreen ratios versus total fluorescence intensity for HeLa cells expressing rFeRepS. The upper left panel (A) shows control cells (grey circles), while the upper right panel (A) displays cells treated with 150 µM FeSO₄ and 250 µM Vitamin C (magenta circles). The lower left panel (A) depicts cells treated with 30 µM ELT (blue circles), and the lower right panel (A) shows cells treated with 400 µg/ml IS (red circles). All cells were treated for 8 hours prior to imaging. **B**) Left: Overlay of control cells (grey circles) and ELT-treated cells (blue circles), showing the distinct shifts in ratio distribution. Right: Overlay of cells treated with FeSO₄ and IS, illustrating differences in ratio distributions following these treatments. **C**) Pie charts illustrating the percentage distribution of ratio values for each treatment condition, as indicated.

Most cells showed a ratio between 0.5 and 5.0, although a subset exhibited ratios below 0.5 or above 5.0, even under control conditions (Figure 7A, left upper panel). Although this system is not yet fully calibrated, we hypothesize that cells with ratios below 0.5 are likely iron-deficient, while those with ratios above 5.0 may be experiencing iron overload. Within the 0.5 to 5.0 range, we also observed considerable heterogeneity, suggesting that while some of these cells may have a balanced iron status, others might already be experiencing either iron deficiency or iron excess.

Next, we explored how individual cells of the cell population responded to iron loading via Fe(II) sulfate and Vitamin C treatment. As shown in **Figure 7A** (**right upper panel, B and C**), this treatment did not markedly alter the ratio distribution across the population. While a modest increase in the number of cells with ratios around 5.0 was observed, cells with extremely high ratios were largely absent, possibly due to the cytotoxicity of excess iron, which may trigger ferroptosis in iron-overloaded cells. Interestingly, the population of cells with ratios below 0.5 remained largely unaffected by Fe(II) treatment, suggesting that certain cells continued to display signs of limited iron uptake despite the increased extracellular iron availability. This observation raises the possibility that HeLa cells may have a limited capacity to take up Fe(II) through the plasma membrane, potentially due to varying levels of DMT1, a key ion channel involved in Fe(II) uptake across the plasma membrane.

To probe the system’s sensitivity to iron depletion, we treated rFeRepS-expressing HeLa cells with ELT and analysed the resulting mCherry/mNeonGreen ratio values plotted against total fluorescence intensity (**Figure 7A and B, lower left panel, and C**). ELT treatment led to a marked reduction in the number of cells with ratio values around or above 5.0, while significantly increasing the proportion of cells with low ratios (**Figure 7A and B, lower left panel, and C**). These findings demonstrate that rFeRepS effectively visualizes shifts toward iron deficiency in response to iron chelation by ELT. Furthermore, these experiments suggest that ELT treatment in HeLa cells may lower ISC synthesis and likely upregulate TfR expression—both signs of cellular iron "hunger" as cancer cells adjust to iron depletion. In contrast, treatment with carbohydrate-based iron sucrose nanoparticles (IS) effectively increased the proportion of cells with a higher mCherry/mNeonGreen ratio around or above 5.0, while simultaneously reducing the number of cells with a low ratio below 0.5 (**Figure 7A, right lower panel, B and C**). Unlike treatment with Fe(II) salts, which had a limited impact on alleviating iron deficiency, IS nanoparticle treatment proved to be more effective at increasing intracellular iron levels in HeLa cells. These results suggest that rFeRepS can distinguish the relative efficacy of iron sources in vitro, indicating that IS nanoparticles are more readily taken up by HeLa cells and lead to a more robust increase in iron availability than Fe (II) salts.

To further validate that the observed changes in mCherry/mNeonGreen ratios were a result of alterations in cellular iron status, we also treated cells expressing the control construct with the same substances. As shown in **Figure 8**, the ratio values remained consistent at around 1 across all treatments, suggesting that the changes in the rFeRepS-expressing cells were specifically related to iron modulation, and not due to any non-specific effects on fluorescence or expression levels.

**Figure 8.**
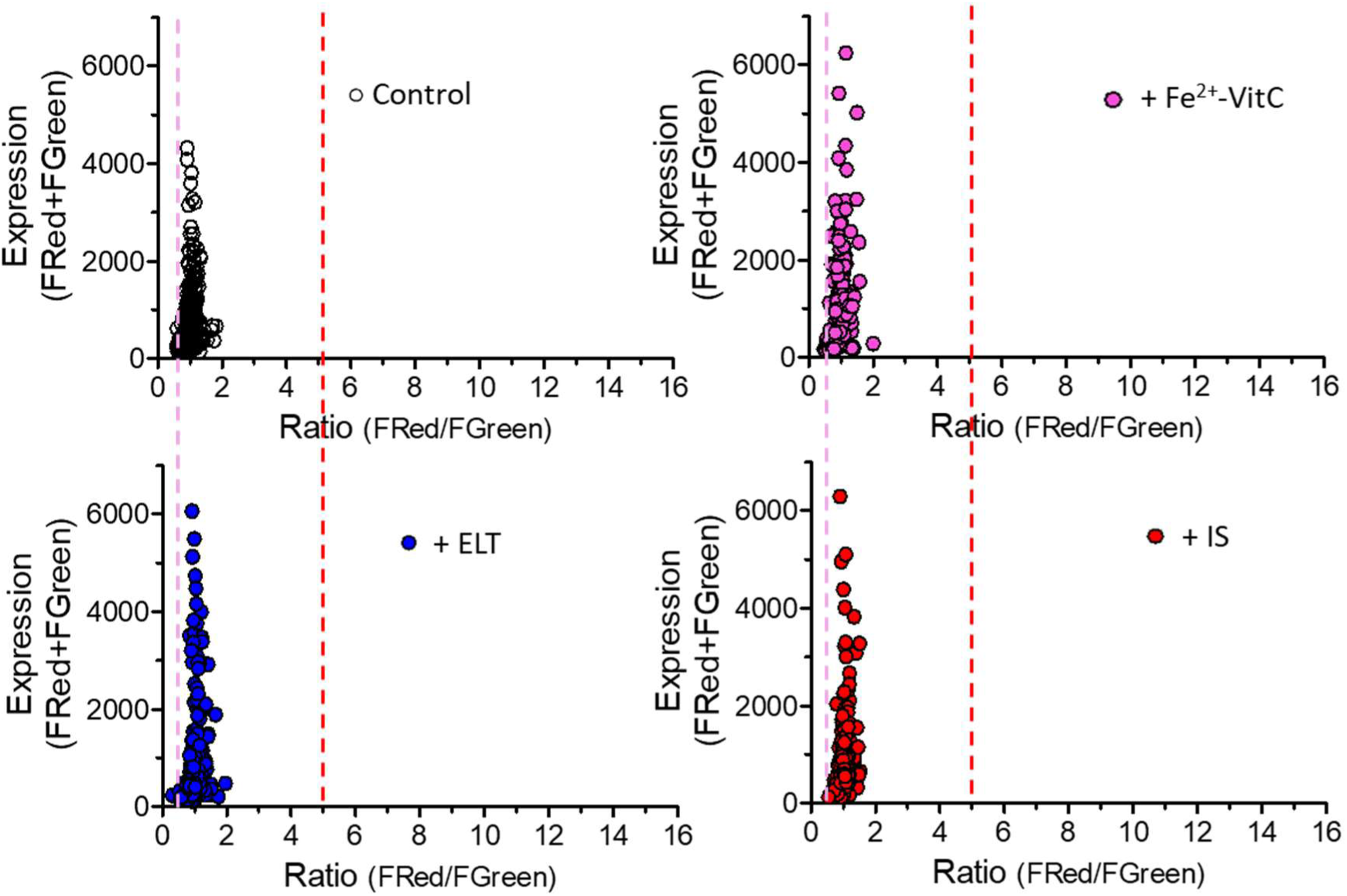
mCherry/mNeonGreen Ratio Distribution in Control Construct-Expressing HeLa Cells Under Different Treatments. XY plots showing the distribution of mCherry/mNeonGreen ratios versus total fluorescence intensity for HeLa cells expressing the control construct. The grey circles represent untreated control cells, magenta circles indicate cells treated with 150 µM FeSO₄ and 250 µM Vitamin C, blue circles show cells treated with 30 µM ELT, and red circles represent cells treated with 400 µg/ml IS. All cells were treated for 8 hours before imaging. The ratio values remain consistent across all treatments.

### IronFist Principle and Design 1: A Functional Prototype for cellular Fe²^+^ sensing

Next, we tested the first prototype of the IronFist system, as illustrated in **Figures 9A** and **D**. This design 1 uses a vector that drives the expression of a construct under the control of the moderate hPGK promoter (**Figure 9 D**), as described above. The construct includes an Hr domain derived from the FBXL5 protein (for details see Figure 2), which is known to regulate iron homeostasis (Figure 2). This design fuses the Hr domain with mTagBFP2 (a blue fluorescent protein). We hypothesized that such a construct would be degraded via the ubiquitin-proteasome system under conditions of low Fe²^+^ availability. However, when the labile Fe²^+^ pool rises, Fe²^+^ ions bind to the Hr domain, stabilizing the fusion protein and preventing its degradation. As a result, the whole construct including mTagBFP2 accumulates and its fluorescence signal becomes detectable. This design provides a dynamic, Fe²^+^-dependent readout of cellular iron levels. A combination of rFeRepS and Iron Fist design to connect LIP and ISC synthesis and assess the interconnection of both has not been performed yet.

**Figure 9.**
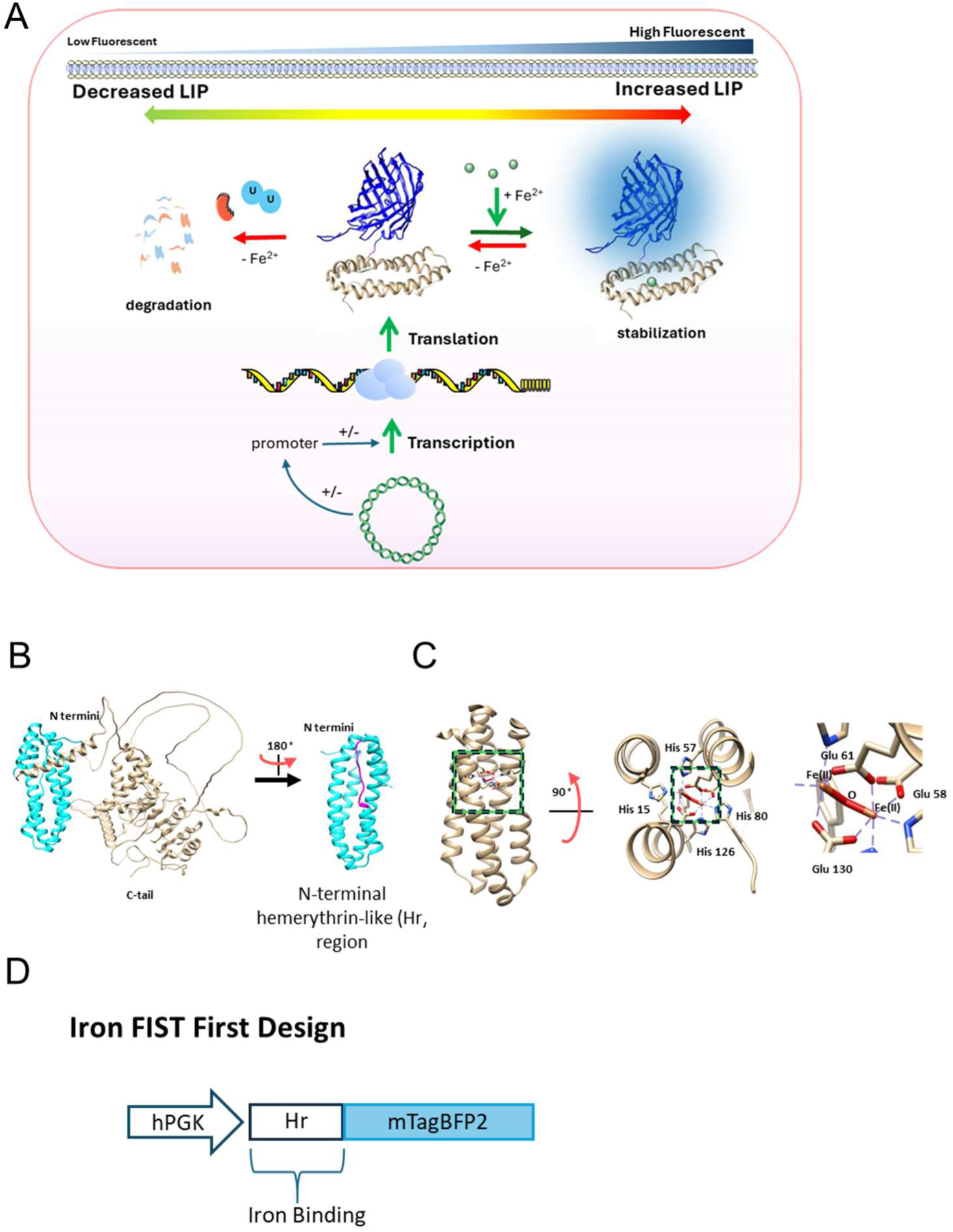
IronFist Design 1: Mechanism and Structural Basis. **A)** Schematic representation of the IronFist Design 1 system, featuring a blue fluorescent protein (mTagBFP2). The vector is transcribed into mRNA, which is then translated into a fusion protein. In the absence of Fe²^+^, the fusion protein is unstable due to ubiquitination and subsequent degradation. However, when Fe²^+^ binds to the Hr domain, the protein stabilizes, preventing degradation and allowing the accumulation of blue fluorescence as a sign of increased labile iron pool (LIP). **B** and **C**) Detailed structural representations of the Hr domain from the FBXL5 protein. The cyan structure highlights the hemerythrin-like domain, and the magenta represents the iron-dependent degron sequence. Zoom-in views show the critical interaction between His 80 and Fe²^+^-O-Fe²^+^, which facilitates protein stabilization in the presence of iron. **D)** Schematic of the DNA sequence encoding the IronFist Design1.

### Structural Basis of the IronFist System

The core mechanism of the IronFist system is based on the iron-regulated degradation of the fusion protein, which is driven by the Hr domain derived from the FBXL5 protein. The predicted structure of FBXL5 and the crystal structure of its Hr domain are shown in **Figure 9B** and **C**. The Hr domain contains a hemerythrin-like structure (cyan) that houses an iron-dependent degron sequence (Chollangi et al., 2012; Ruiz & Bruick, 2014) (magenta). This sequence, located within amino acids 75-85, is critical for regulating the stability of the protein in response to iron levels (Ruiz et al., 2013; Thompson et al., 2012; Wang et al., 2020). Zoom-ins at different angles in **Figure 9B** and **C** illustrate the key amino acid residues involved in iron coordination. Specifically, His 80, located within the degron region, plays a crucial role in the interaction with the Fe²^+^-O-Fe²^+^ complex. In the absence of Fe²^+^, the degron is exposed, leading to ubiquitination and degradation of the protein (Thompson et al., 2012)However, when Fe²^+^ binds to the His 80 residue, it stabilizes the protein by preventing the degron from being recognized by the degradation machinery, thus preventing its ubiquitination and degradation(Ruiz & Bruick, 2014). This structure-function relationship underpins the IronFist system, allowing for a dynamic and Fe²^+^-sensitive response that can be used to monitor cellular iron levels.

### Functional evaluation of IronFist Design 1 in HeLa cells

To assess the functionality of the IronFist Design 1 construct, we co-transfected HeLa cells with the IronFist construct and mCherry, a red fluorescent protein, and selected cells that exhibited both red and faint blue fluorescence. The presence of both fluorescence signals indicated expression of the IronFist construct. Subsequently, we treated the cells with Fe (II) sulfate and Vitamin C to elevate iron levels and monitored fluorescence signals for both channels over time using live cell imaging.

Upon treatment with Fe²⁺, we observed a significant increase in blue fluorescence over time in many cells, while red fluorescence remained constant (**Figure 10 A**). This result demonstrated that the IronFist system functions as intended, with an increase in labile iron pool (LIP) triggering the stabilization of the IronFist construct and a corresponding increase in blue fluorescence.

**Figure 10:**
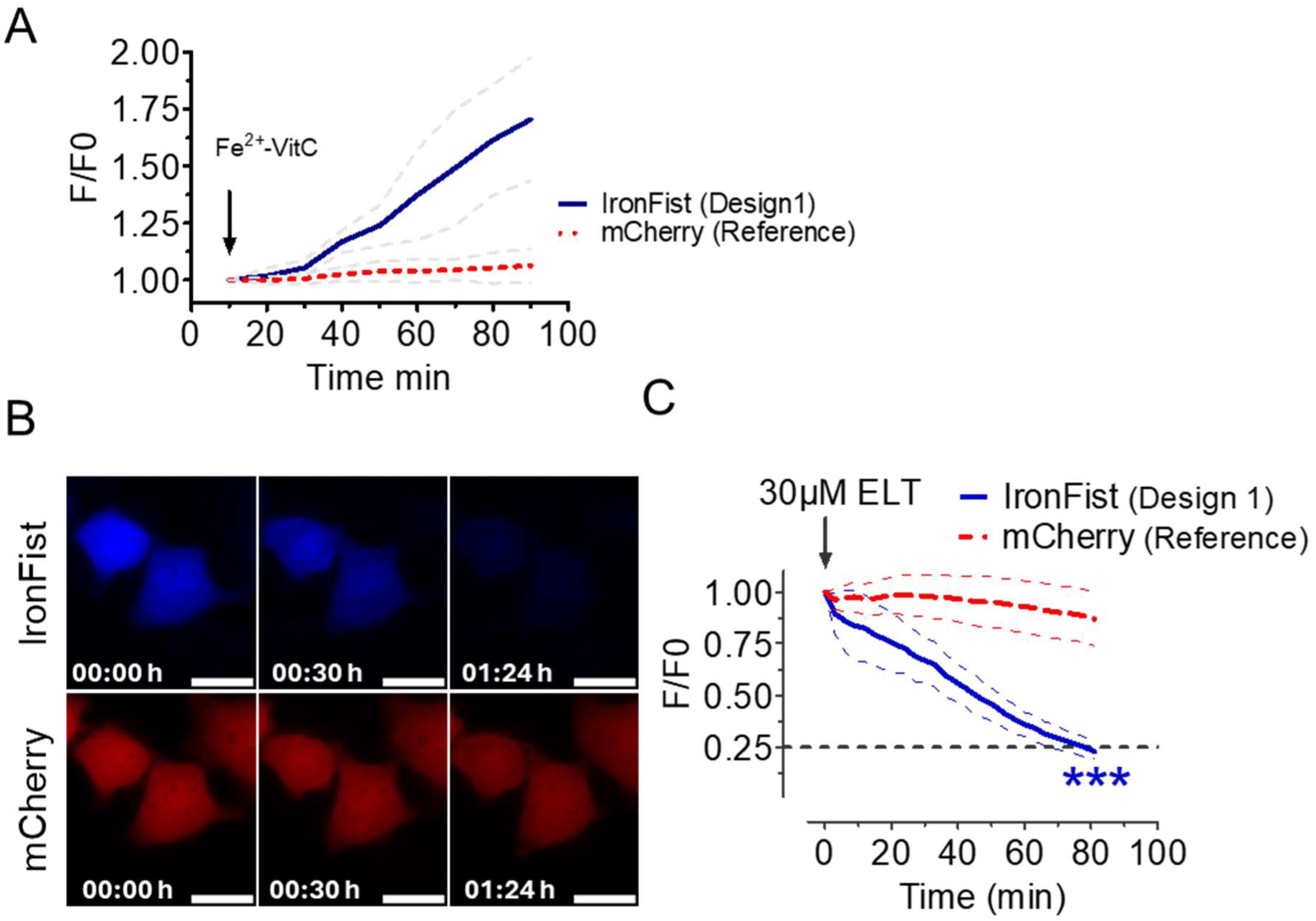
Functional Evaluation and Dynamic Response of IronFist Design 1 in HeLa Cells. **A**) Curve graph showing the average intensity ± SD (n=11) of blue fluorescence from mTagBFP2 (IronFist construct) and mCherry over time in response to Fe²⁺ addition. HeLa cells were co-transfected with IronFist Design1 and mCherry; cells expressing both constructs were selected. **B)** Representative images of HeLa cells pre-treated with FeSO_4_/Vit C for 2 hours and expressing IronFist Design 1 together with mCherry before and during treatment with the iron chelator ELT. The images show the decrease in blue fluorescence (IronFist) over time, indicating the response of the construct to iron depletion. **C)** Corresponding intensity curves for the blue fluorescence (IronFist) and red fluorescence (mCherry, iron-insensitive reference protein) during the same treatment as shown in Panel B. *** indicate significant differences between mCherry and mTagBFP2 fluorescence intensity at given time points. p<0.0001.

To further assess the dynamic sensitivity of IronFist Design 1 to cellular iron levels, we first treated HeLa cells with Fe²⁺ to increase their iron content and then added ELT (iron chelator) during time-lapse imaging. While the red fluorescence from the iron-sensitive mCherry remained stable, the blue fluorescence from the IronFist construct gradually decreased over time (**Figure 10 B** and **C**). This observation indicates that IronFist Design 1 is capable of dynamically and reversibly reporting changes in cellular iron levels, with a decrease in blue fluorescence corresponding to the reduction of the labile iron pool (LIP) upon iron chelation by ELT. This confirms the responsiveness of the system to both iron supplementation and depletion, further validating its potential as a tool for monitoring cellular iron dynamics.

### Functional evaluation of IronFist Design 2: a ratio metric tool

To further improve the ratio metric sensing of the labile iron pool (LIP), we developed IronFist Design 2, which incorporates a green/red fluorescence ratio system. In this design, the Hr domain is fused with mNeonGreen, a bright green fluorescent protein variant, while mCherry, an iron-insensitive reference protein, is expressed via a ribosomal skipping sequence (**Figure 11 A** and **B**). This configuration enables the ratio metric detection of cellular iron status by measuring the green (mNeonGreen) and red (mCherry) fluorescence intensities.

**Figure 11:**
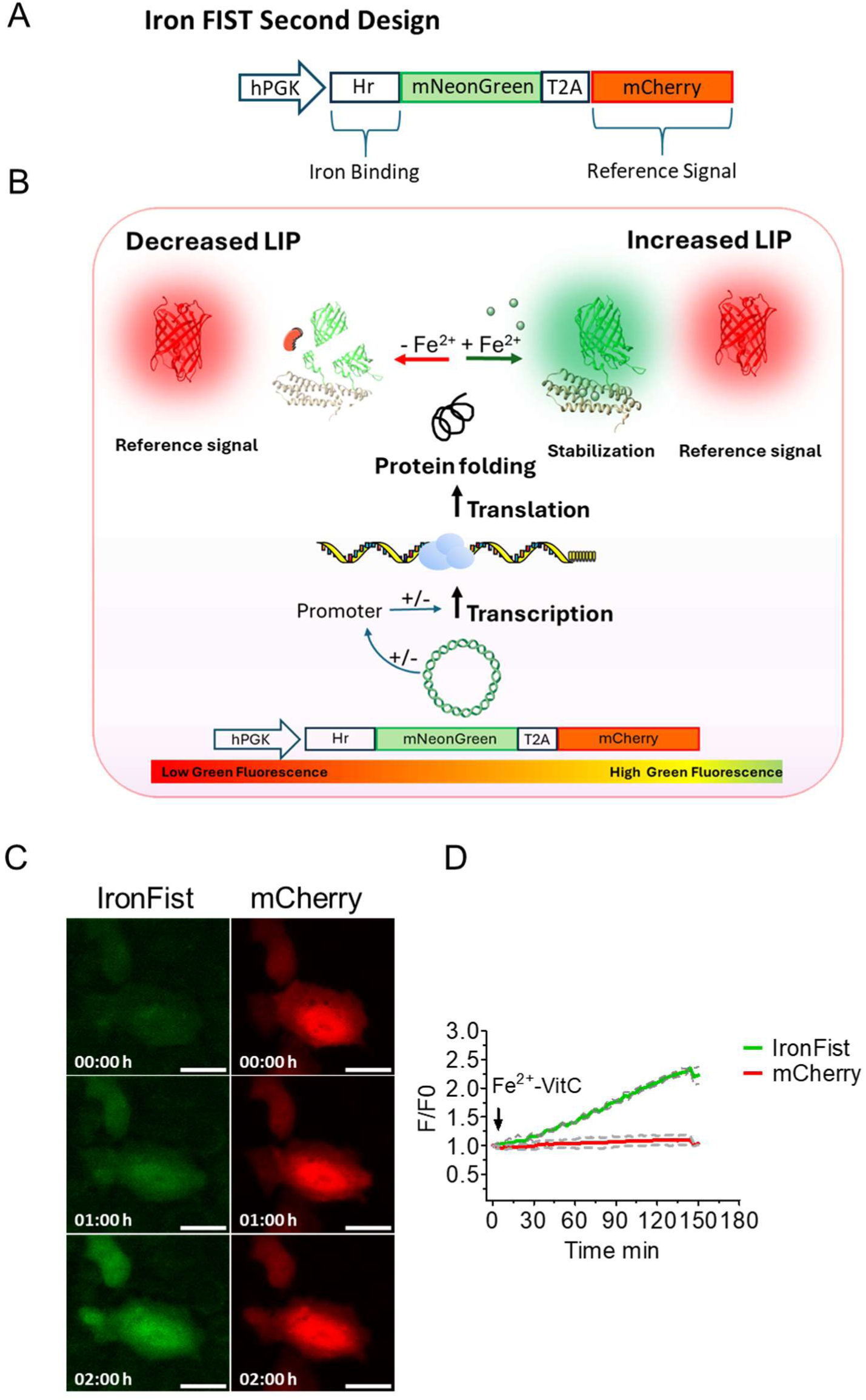
Functional Evaluation of IronFist Design 2: A Ratiometric Tool for Monitoring the Labile Iron Pool (LIP) **A)** Schematic representation of the DNA design for IronFist Design 2, incorporating a Hr domain fused to mNeonGreen (green fluorescent protein) and mCherry (red fluorescent protein). The mCherry is expressed via a ribosomal skipping sequence, serving as an iron-insensitive reference protein. The Hr-mNeonGreen fusion responds to changes in cellular iron levels, allowing for ratiometric detection of the LIP. **B)** Cartoon illustrating the principle of IronFist Design 2. Upon binding of Fe²⁺ to the Hr domain, the green fluorescence (mNeonGreen) increases, while the red fluorescence (mCherry) remains unaffected, providing a ratiometric signal of the LIP. **C)** Representative time-lapse images of HeLa cells expressing IronFist Design 2 before and after treatment with FeSO₄ and Vitamin C. The green fluorescence (mNeonGreen) increases following iron treatment, while the red fluorescence (mCherry) remains constant, indicating the system’s sensitivity to iron fluctuations and its potential for monitoring the LIP. **D)** Time-lapse analysis of the green/red fluorescence ratio in HeLa cells expressing IronFist Design 2. The green curve indicates the increase in green fluorescence (mNeonGreen) over time, reflecting the increase in the LIP, while the red curve shows the stable red fluorescence (mCherry), confirming the ratiometric functionality of the system.

To test the functionality of IronFist Design 2, we examined its performance in HeLa cells. Upon treatment with FeSO_4_/Vit C, we observed a significant increase in the green/red fluorescence ratio, as shown in **Figure 11C** and **Figure 11D**. These data demonstrate that IronFist Design 2 is sensitive to changes in cellular iron levels and can effectively monitor the LIP through ratio metric fluorescence time-lapse imaging

### End-point analysis of IronFist ratiometric response in HeLa Cells

To evaluate the response of IronFist Design 2 compared to a control construct, we measured the mNeonGreen/mCherry fluorescence ratio in HeLa cells under various conditions. HeLa cells were transfected with either IronFist or the iron-insensitive control construct and imaged two days post-transfection. Under untreated conditions, cells expressing IronFist exhibited a significantly lower mNeonGreen/mCherry ratio (0.07) than those expressing the control construct (1.03), indicating insufficient labile iron pool (LIP) to stabilize IronFist in the absence of supplemental iron (**Figure 12A**). To confirm that the Hr-mNeonGreen component of IronFist undergoes proteasomal degradation, cells expressing IronFist were treated with the proteasome inhibitor MG132 for 4 and 6 hours. This treatment increased the mNeonGreen/mCherry ratio by selectively enhancing green fluorescence intensity, validating that Hr-mNeonGreen degradation occurs through the proteasomal pathway (**Figure 12B**).

**Figure 12:**
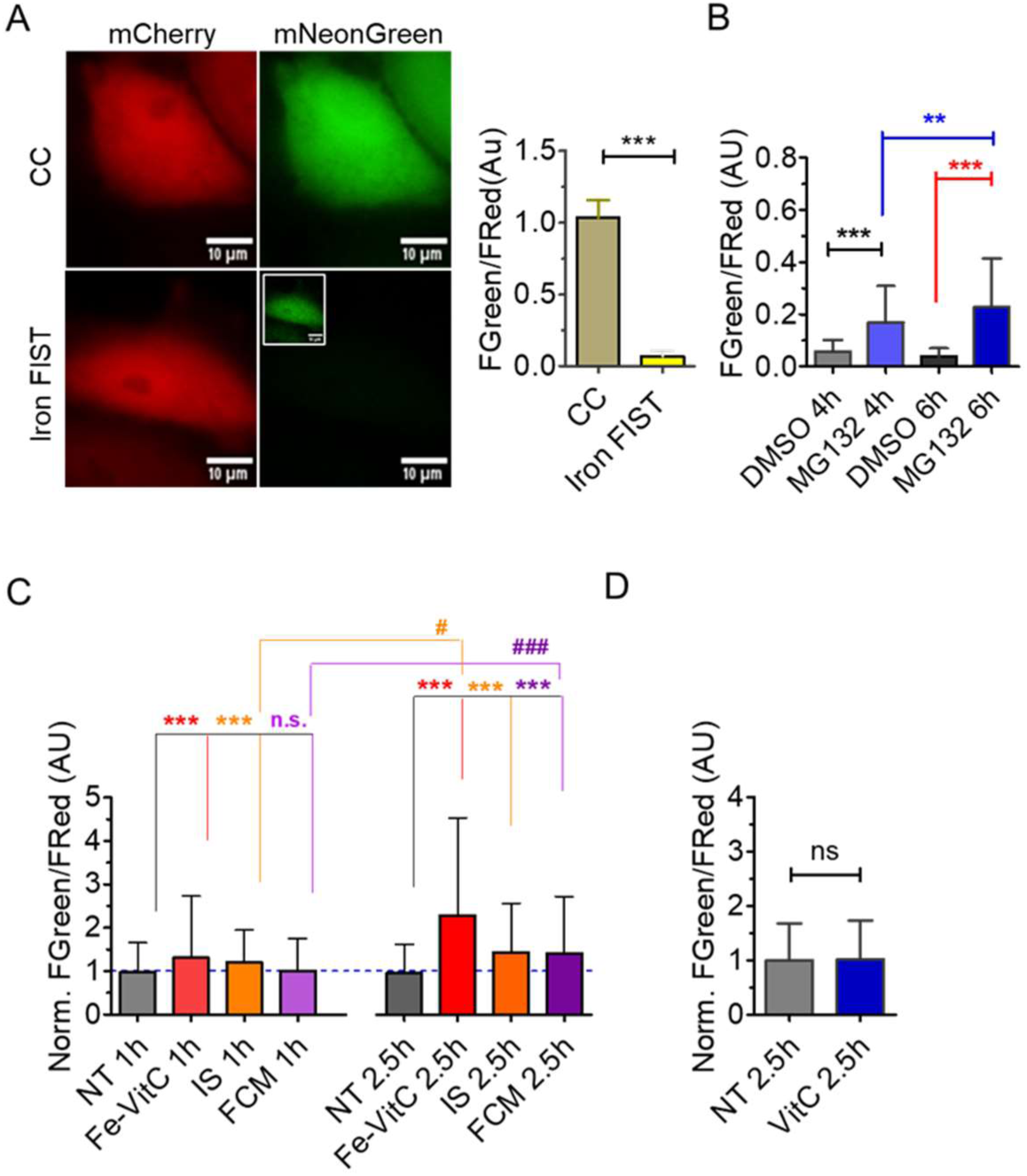
Differential Response of the IronFist to Iron, Vitamin C, and Proteasome Inhibition in HeLa Cells. **A**) Representative images of HeLa cells expressing IronFist and the control construct, expression of mNeonGreen in ironFist treated cell shown in smaller brightness contrast setting in inset image, with corresponding mNeonGreen/mCherry ratio plots. In the ratio plot (right panel), the dark yellow column represents the control construct, and the yellow column represents IronFist. Under non-iron-treated conditions. **B**) mNeonGreen/mCherry ratio plots of HeLa cells expressing IronFist following treatment with 10 µM MG132 for 4 hours (blue) and 6 hours (dark blue). Treatment for n of DMSO 4h =3wells/114 cells n of DMSO 6 hours=3wells/118 cells no f MG132 4h=3wells/90 cells. n of MG132 6h=3wells/129cells.**C**) Normalized mNeonGreen/mCherry ratio plots of HeLa cells expressing IronFist treated with either 150 µM FeSO₄ and 250 µM Vitamin C (red), 500 µg/ml IS (orange), or 500 µg/ml FCM (purple) for 1 and 2.5 hours. FeSO₄-Vitamin C treatment. n of 1h NT= 9wells/240 cells, n of 1h FeSo_4_-VitC= 8wells/185 cells, n of 1h IS= 7wells/179 cells, n of 1h FCM= 9 wells/269 cells 2,5h NT= 9wells/240 cells, n of 2,5h Fe-VitC= 9wells/248 cells, n of 2,5h IS= 9wells/313 cells, n of 2,5h FCM= 9wells/256 cells. **D**) Normalized mNeonGreen/mCherry ratio of HeLa cells expressing IronFist treated with 250 µM Vitamin C (blue) compared to untreated cells (dark grey). n of NT= 9wells/244 cells, n of VitC= 7wells/160 cells. DATA=mean ±SD, statistical analysis non-parametric t test, *P<0.05, **P<0.001 ***P<0.0001 statistically significant.

Further testing involved treating HeLa cells with FeSO₄/Vitamin C, iron supplementation (IS), or ferric carboxymaltose (FCM) for 1 and 2.5 hours, followed by end-point imaging. FeSO₄/Vitamin C treatment produced a clear increase in the mNeonGreen/mCherry ratio at 1 hour, with a more pronounced effect observed at 2.5 hours (**Figure 12C**). Similar results were also obtained from Hela cells stably expressing Iron Fist design 2 with 2 hours of FeSO₄/Vitamin C treatment (data not shown). IS treatment produced only a slight increase in the ratio at 1 hour, but the effect became more evident by 2.5 hours. In contrast, FCM did not affect the ratio at 1 hour but increased it by 2.5 hours, indicating a slower response relative to IS and FeSO₄/Vitamin C. To ascertain whether Vitamin C alone influences the IronFist ratio, cells were treated with Vitamin C without iron supplementation. Vitamin C alone did not alter the mNeonGreen/mCherry ratio significantly at 2.5 hours, suggesting that the observed ratio changes were specific to iron exposure in the FeSO₄/Vitamin C condition **(Figure 12D**).

### Broad detection window from 24 to 42 hours post-transfection for reliable iron sensing with IronFist Design 2

We hypothesized that following transfection, mRNA and expression levels of the IronFist construct would progressively increase, reaching a peak due to rising rates in transcription, translation, and protein folding before gradually declining due to cell division and probably other processes. Consequently, the timing post-transfection could potentially influence the ratiometric output of the IronFist Design 2 system. However, our findings showed that at both 24- and 42-hours post-transfection, FeSO₄/Vitamin C treatment significantly increased mNeonGreen fluorescence without affecting the iron-insensitive mCherry reference fluorescence, leading to a marked increase in the green/red ratio at both time points (**Figure 13 A and B**). As expected, mCherry fluorescence intensity was significantly higher at 42 hours compared to 24 hours (**Figure 13 A, left upper panel right**), reflecting the accumulation of this protein over time in cells expressing the construct. However, mNeonGreen fluorescence remained comparably low at both time points before iron treatment, increasing significantly only upon FeSO₄/Vitamin C addition (**Figure 13 A, left upper panel left**).

**Figure 13.**
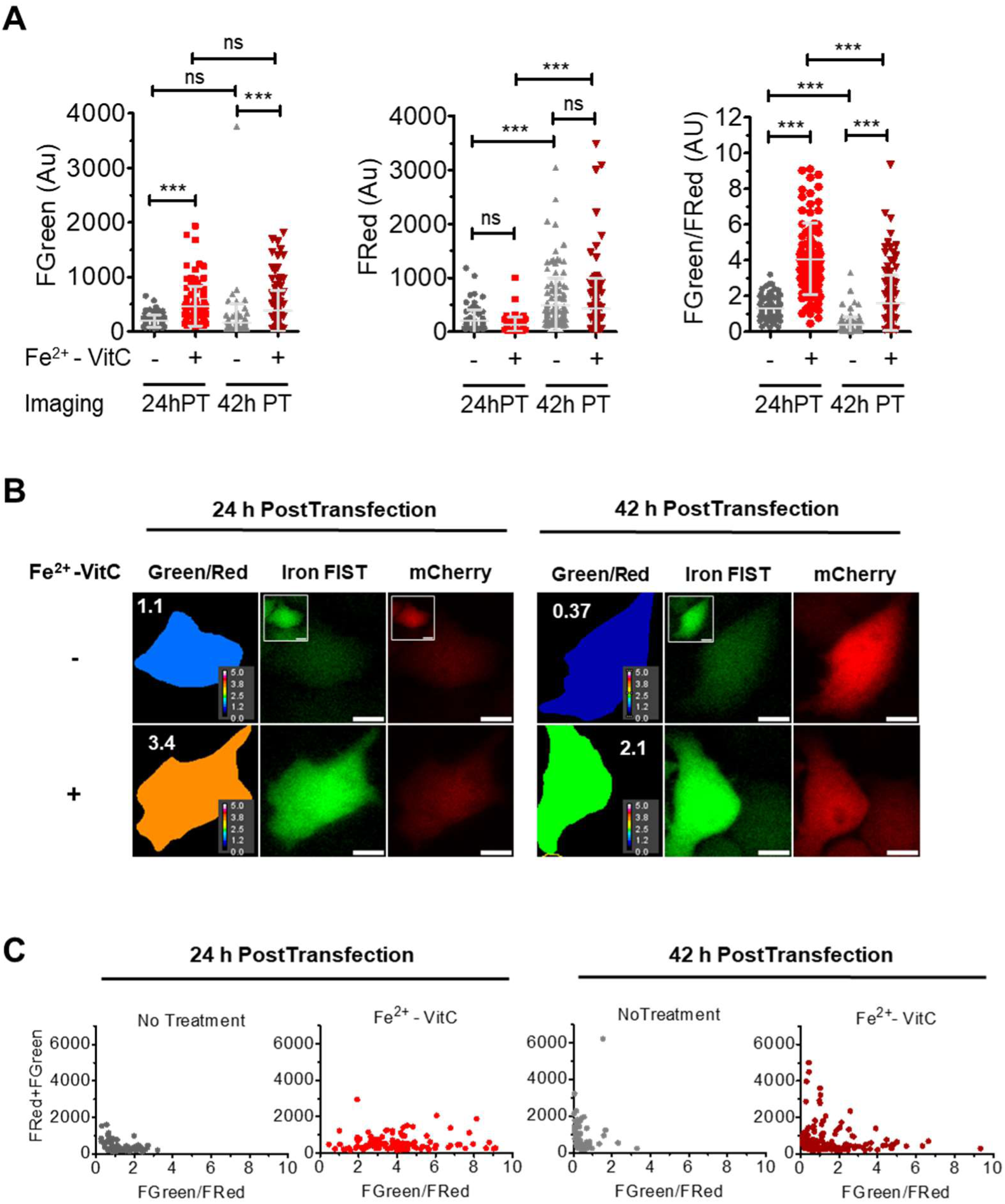
IronFist Design 2 functionality over time post-transfection. **A**) Bar graphs showing fluorescence intensities and mNeonGreen/mCherry ratios at 24 hours (n=80 for NT, grey column; n=94 for Fe/VitC treated, red column) and for 42 hours (n=80 for NT, grey column; n=132 for Fe/VitC treated, red column) post-transfection, following treatment with FeSO₄ (150 µM) and Vitamin C (250 µM) for 2.5 h. In the top-left panel, average mNeonGreen intensities are displayed: the first grey column represents untreated cells (NT, n=80) at 24 hours post-transfection, followed by the second red column for treated cells at the same time point (n=94). The next two columns show data for 42 hours post-transfection, with grey for untreated cells (NT, n=125)) and red for treated cells (n=132). The top-right panel displays corresponding values for the mCherry reference protein. In the lower panel, mNeonGreen/mCherry ratios for both time points are shown, comparing untreated (grey) and treated cells (red) to illustrate changes in iron sensitivity. DATA=mean ±SD. ns indicates non-significant and *** significant differences P<0.001. **B**) Representative images of HeLa cells transfected with IronFist Design 2, with treatment conducted 24- or 42-hours post-transfection. The left set shows untreated and treated cells imaged 24 hours post-transfection, and the right set shows images taken at 42 hours. Each set displays the generated ratio images (top row), mNeonGreen fluorescence (middle row), and mCherry fluorescence (bottom row). **C**) Scatter plots display the mNeonGreen + mCherry sum (Y) plotted against the mNeonGreen/mCherry ratio values (X) of all the individual cells at both 24 (left) and 42 (right) hours, highlighting the absence of correlation between expression levels and ratio

Additionally, we found no correlation between the mNeonGreen/mCherry ratio and the absolute expression levels, regardless of iron treatment (**Figure 13 C**, upper panels for untreated; lower panels for FeSO₄/Vitamin C treated). This suggests that the ratiometric sensitivity of IronFist is reliable across a range of expression levels and times post-transfection.

Overall, these experiments demonstrate that transient transfection of IronFist provides a substantial operational window, from at least 24 to 42 hours post-transfection, within which the system remains responsive to cellular iron changes and effectively visualizes alterations in the labile iron pool (LIP).

### Long-term monitoring of IronFist design 2 reveals heterogeneous LIP responses in single cells

We conducted long-term live-cell imaging to monitor the green/red ratiometric IronFist Design 2 in individual HeLa cells over an 8-hour period, capturing images every 3 minutes for both mNeonGreen and mCherry fluorescence channels. Just before initiating this time-lapse experiment, cells were treated with FeSO₄ and Vitamin C to elevate the labile iron pool (LIP). **Figure 14** presents three distinct response patterns to highlight the heterogeneity of cellular reactions to iron treatment:

**Figure 14:**
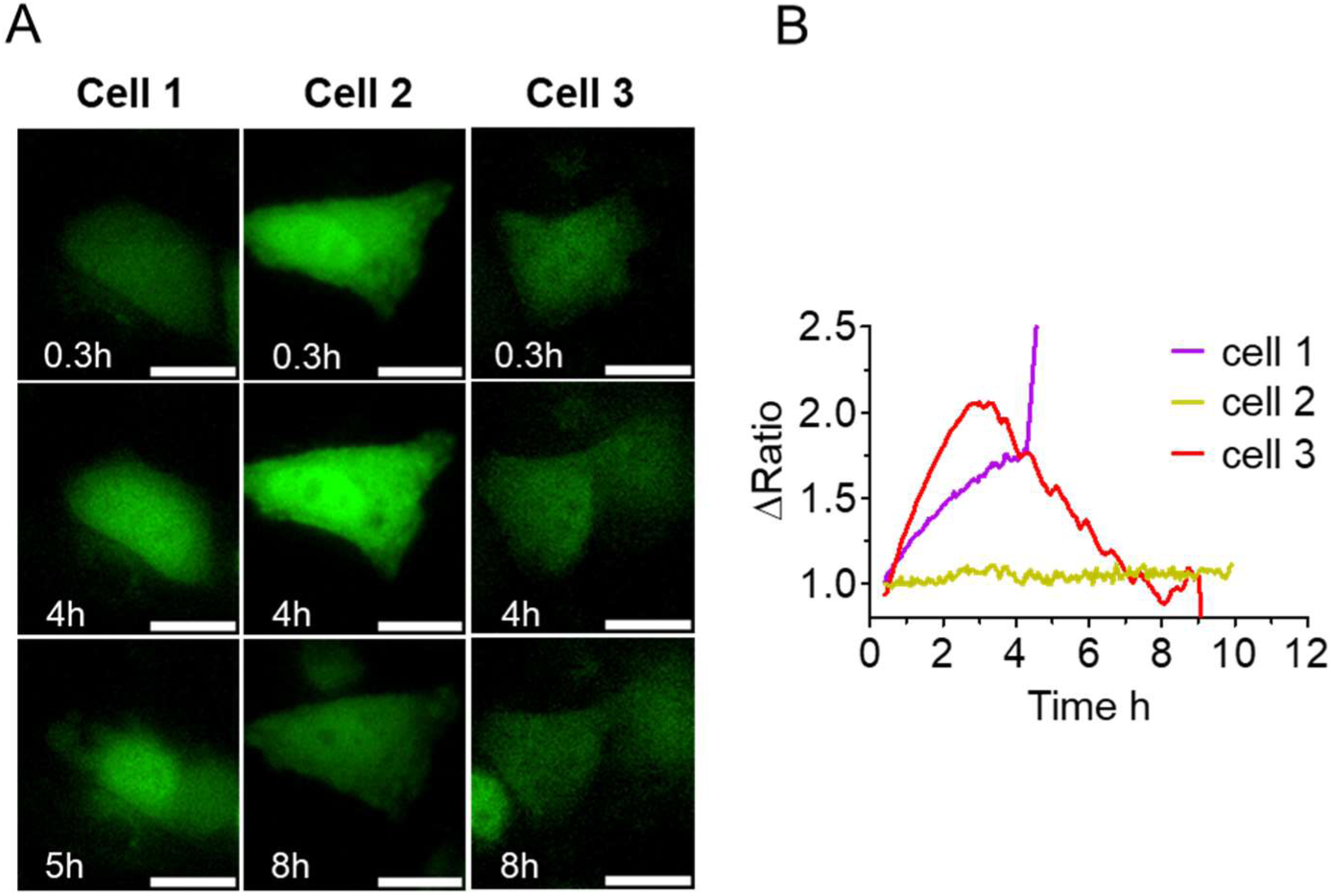
Single-cell analysis of LIP dynamics using IronFist design 2. **A**) Representative time-lapse images of three HeLa cells, each responding differently to FeSO₄ (150 µM) and Vitamin C (250 µM) treatment, demonstrating variability in labile iron pool (LIP) dynamics. Green (mNeonGreen) fluorescence channels are shown for each cell at three selected time points across the 8-hour monitoring period as indicated. **B)** Normalized ratio curves showing the green/red fluorescence ratio over time for each cell, highlighting distinct LIP responses. Ratio values were normalized to the initial ratio at the start of the measurement to emphasize changes. Cell 1 (violet trace) shows a gradual increase in LIP over approximately 4 hours followed by cell death, likely due to iron toxicity. Cell 2 (yellow-green trace) maintains a stable ratio over the entire period, indicating no significant LIP increase. Cell 3 (red trace) shows an initial increase in the ratio, followed by a return to baseline, suggesting an initial LIP rise that the cell subsequently manages to regulate.

In *Cell 1*, an immediate, continuous increase in the green/red fluorescence ratio was observed following Fe(II) treatment, suggesting a progressive buildup of the LIP over 4 hours. After this period, the cell likely was affected by iron toxicity and died, indicated by an abrupt cessation of further ratio changes. *Cell 2* exhibited an initial increase in the green/red ratio upon iron addition, followed by a gradual return to baseline levels after about 3 hours. This cell appeared to respond by first increasing the LIP, then successfully regulating and reducing it back to starting levels. Finally, *Cell 3*, by contrast, displayed no significant change in the green/red ratio throughout the experiment and remained viable, suggesting that Fe(II) treatment did not induce a substantial increase in the LIP for this cell.

These results demonstrate the utility of the ratiometric IronFist system for visualizing dynamic, cell-specific fluctuations in the LIP, enabling real-time insights into individual cellular responses to iron treatment.

## Acknowledgments

This study was supported by the Medical University of Graz, Austria and CSL Vifor, Switzerland. IS and FCM nanoparticles were a kind gift from CSL Vifor. Special thanks to our colleagues from the Medical University of Graz, Yusuf Ceyhun Erdogan, and Anna Lischnig for their discussions about cellular iron dynamics. We thank Anna Schreilechner, Dr. Rene Rost, and Stella Petritsch for technical assistance, cell culture, and preparation of material.

## Author Contributions

A.A. conceptualized the study, designed and performed experiments, and analyzed data, A.A. and R.M. wrote the manuscript. A.A., R.M., and W.F.G. contributed to experimental design, data analysis, and manuscript editing. Ş.Ç. and E.E. generated Hela cells stably expressing IronFist performed FACS experiments and analysed the data. A.B.A., R.D., and B.F. provided materials and knowledge and helped with the experimental designs. B.G. wrote the Fiji macro script for image analysis. R.M., E.E., W.F.G., and B.F. supervised the study, provided guidance on experimental approaches, and critically revised the manuscript. All authors reviewed and approved the final manuscript.

## Materials and Methods

### Buffers and Solutions

Cell culture materials were obtained from Grainer Bio-One (Kremsmünster, Austria). Experimental buffer 1 prepared in house with following ingredients and concentrations (in mM); 2 CaCl_2_, 135 NaCl, 1 MgCl_2_, 5 KCl, 10 HEPES, 2.6 NaHCO_3_, 0.44 KH_2_PO_4_, 0.34 Na_2_HPO_4_, 10 D-Glucose, 2 L-Glutamine and supplemented with 1x amino acids, 1x vitamins (pH=7.45)

### Compounds

Eltrombopag (ELT), MG 132, Erastin-2 (Ers-2) were purchased from MedChem Express (#HY-15306, #HY-13259, #HY-139087, respectively) purchased from MedChem Express and dissolved in DMSO at 10 mM concentration then aliquoted and kept at -20 °C for further use and used at 10 µM final concentration. IS np, (ferric iron sucrose nanoparticle) and FCM (Ferric carboxy maltose) were kind gifts from Vifor pharma. Ferrous sulphate heptahydrate was purchased from Sigma Aldrich #215422. Ascorbic acid is obtained from Aponorm. Fe-Vitamin C mix prepared freshly by dissolving both compounds in double distilled water. NucBlue™ was purchased from ThermoFisher Scientific #33342.

### Molecular Cloning and design

#### IronFists

First prototype of iron FIST designed by fusing a blue fluorescent protein, mtagBFP2 to Hr domain of FBXL5. To generate pseudo ratio metric tool, we have designed a vector which has a fusion of Hr domain to a very bright version of GFP (mNeonGreen) followed by a ribosomal skipping sequence, T2A, and mCherry fluorescent protein as a reference signal. As non-iron dependent control we also designed a control vector which expresses both fluorescent proteins under the same promoter. We have selected a moderate strength promoter, hPGK (human phosphoglycerate kinase) to not overwhelm the proteasomal degradation note that we have not yet tested other promoters. All three vectors have been synthetized by Vector Builder (Neu-Isenburg, Germany).

#### FeRepS

Plasmids encoding LoFeRepG1, HiFeRepR1, Hi FeRep R1-IRE and, mCh_T2A_mNeonGreen (control construct) carrying plasmids synthesized and cloned by Vector Builder (Neu-Isenburg, Germany). PcDNA 3.1-encoding BgH poly A tail signal, T7 terminator, and rrnG terminator between EcoRI and BamHI restriction sites (here and on referred as PAT&T in PcDNA3.1-), was purchased from Gene Universal (Delaware, USA). To express both FeReps in one vector HiFeRepR1 was PCR amplified with forward primer carrying BglII enzyme site in the 5’ and EcoRI restriction site at the 3’ end then ligated into PAT&T in PcDNA 3.1-. After confirming the insertion of HiFeRepR1 into vector, LoFeRepG1 and HiFeRepR1 carrying PAT&T PcDNA 3.1-plasmids were cut by BamHI and HinDIII enzymes and then ligated. The resulting vector carrying both FeReps was named rFeRepS.

### Cell Culture and Transfection

Hela S3 cells were cultured in Dulbecco’s modified Eagle medium (DMEM D5523, Sigma Aldrich) supplemented with 10% FCS, 1% penicillin-streptomycin, 1.25mg/mL amphotericin B and 25 mM HEPES (pH=7.45). Cells were routinely passaged every 2-3 days and kept in a humidified cell culture incubator (37°C, 5% CO_2_). Before experiment, cells were seeded on 30mm glass coverslips (Co. KG, Lauda-Königshofen) in 6 well plates (Paul Marienfeld GmbH, Germany). One day after seeding cells were transiently transfected with respective plasmids with Polyjet (Signagen Laboratories, Rockville, USA) according to the manufacturer’s manual. For transfection of a single plasmid 2 µL, and for co-transfection of two plasmids 3 µL of the reagent was used. Transfected cells imaged after 36-42 hours.

### End Point Live Cell Imaging

Transfected Hela S3 cells imaged with Nikon Eclipse Ti2 (Nikon, Austria) equipped with CoolLED pE800 light source (CoolLED, UK, Andover), LED light source with 365, 400, 435, 470, 500, 550, 580, 635, and 740 nm (LEDs is introduced at the back focal plane), Apo 40X/1.15NA water immersion objective, two back-illuminated Kinetix Scientific CMOS cameras (TELEDYNE PHOTOMETRICS, USA, Tucson) mounted to the Nipkow based Crest optics X Light V3 with 70µm pinhole spinning disc system (Crest optics, Italy, Rome) in EPI fluorescent mode by bypassing the spinning disc. The Nikon software (NIS-Elements AR 5.42.06 (Build 1821) LO 64bit, Nikon, Austria) was used for microscope control and image acquisition.

Imaging of Hela cells expressing LoFeRep was performed with co staining of nucleus for normalization of positive green cells to cell number. For this, 2 drops of NucBlue™ were added on each well and plate was kept in cell culture incubator for 15 minutes before imaging.

For imaging of Hela cells expressing rFeRepS cells were either treated with IS (400µg iron/mL/35mm dish), Fe^2+^ -Vitamin C mixture, (150µM, 250µM, respectively) or ELT (30µM) for 8 hours or non-treated (control) before imaging. After treatment is completed, cell media aspirated and replaced with EHL solution.

Imaging of Hela cells expressing IronFist, cells were treated with IS and, FCM (500µg iron/mL/35mm dish), or FeSO_4_ -Vitamin C mixture (150µM, 250µM, respectively) for 1 and 2,5 hours After treatment is completed, cell media aspirated and replaced with EHL solution then imaged.

### Kinetic measurement of IronFist using fluorescence microscopy

Dynamic measurements of IronFist (First prototype) cells were either pretreated with Fe^2+^ -Vitamin C mixture for 2 hours or non-pre-treated, then imaging started with FeSO_4_-Vitamin C mixture treatment (150µM, 250µM, respectively) or ELT (30µM) and imaged nearly 2 hours with 3 minutes of interval.

### Time-lapse imaging in the cell culture incubator

Kinetic measurement of Hi FeRep R1, and Hi FeRepR1-IRE (iron insensitive control construct) was performed in 12 well plates and imaged for 2 days in a device, that placed in a cell culture incubator, CellCyteX™ (Cytena, Freiburg Germany) with 10X dry objective with 1 hour interval. First cells were seeded on a plastic clear bottom 12 well plate (Greiner Bio-one, #665180) the next day cells were transfected with Hi FeRep R1, and Hi FeRep R1-IRE carrying plasmids. After overnight incubation, the media was aspirated and changed with 100 µMD DFO (Deferoxamine) or Fe^2+^-VitC (150, 250 µM, respectively), containing fresh medium then the plates were transferred to the imaging device and imaging started with bright field and red channel.

### Image Analysis

Images processed by Fiji software. For two channel intensity calculations a macro script was used. First, the background subtracted by using a rolling ball function. ROIs (regions of interest) were manually drawn around the cells and intensities obtained then transferred to Excel files. Finally, Ratio of and sum of both channels (FRed/FGreen, FGreen/Fred, and FRed+FGreen, where applicable) calculated in Excel. Ratio images were generated by using each cell’s Roi and ratio assigned by the math function of Fiji.

### Data Analysis

The data obtained from microscope images transferred and analysed in GraphPad Prism 5 Software (GraphPad Software, Inc., La Jolla, CA, USA)

